# Social Tolerance and Innovation in Capuchins: socially more tolerant brown capuchins are better problem-solvers than less tolerant white-faced capuchins

**DOI:** 10.1101/2025.09.05.674457

**Authors:** Sandro Sehner, Claudia Fichtel, Peter M. Kappeler, Helene Meunier

## Abstract

Innovativeness and social learning are the pillars of cultural evolution. While the role of social tolerance in social learning has long been acknowledged, its impact on innovativeness remains poorly understood. Here, we test six groups of captive white-faced capuchins (*Cebus capucinus*) and brown capuchins (*Sapajus apella*) to explore and compare the relationship between social tolerance and problem-solving propensities. White-faced capuchins are renowned for their rich repertoire of social behaviours, whereas brown capuchins are known for their diverse foraging repertoire, resulting in a broad range of culturally transmitted behaviours in both species. We performed a co-feeding experiment to assess species differences in social tolerance and a set of open diffusion novel food puzzles to test innovativeness. We measured associations for each possible dyad within each group during both experiments and compared the resulting social networks between conditions. We also measured neophobia in an independent experiment presenting groups with three novel objects. Overall, during both experiments, brown capuchins were more tolerant, and proportionally more individuals spent their time within the testing areas at any given moment. We also found that networks across experimental contexts were more similar in brown capuchins compared to white-faced capuchins. Moreover, although differences in approaching and exploring food puzzles were marginal, the success rate was higher in brown capuchins than in white-faced capuchins. Finally, neither eigenvector centrality within the networks nor neophobia could explain individual problem-solving success. Age was the strongest predictor for successfully extracting a food item. The results indicate that species-level differences in social tolerance may contribute to divergent innovation patterns, whereas individual-level variation within species may have only a minor influence. We conclude that the higher social tolerance observed in brown capuchins during feeding and foraging contexts can contribute to their broader repertoire of foraging behaviours and foraging traditions.

**Data availability:** All data needed to evaluate the conclusions are present at OSF (https://doi.org/10.17605/OSF.IO/BY5MJ).

**Declaration of Interest:** The authors declare no competing interests.

**Author Contributions:** **Sandro Sehner: Writing** – original draft, review & editing, Conceptualization, Methodology, Formal analysis, Data curation, Funding acquisition. **Claudia Fichtel: Writing** – review & editing. **Peter Kappeler: Writing** – review & editing. **Helene Meunier: Writing** - review & editing, Conceptualization, Methodology.

## Introduction

Social tolerance is widely considered a prerequisite for group living as it supports the emergence of social behaviours that mediate both agonistic and affiliative interactions (De Waal, 1986; van de Waal et al., 2010). Accordingly, social tolerance has been defined as “the willingness of individuals to interact non-agonistically with each other and to spend time in proximity to their social partner” (Rina Evasoa et al., 2019). It has been identified as a key mechanism shaping multiple aspects of social life, including cooperation (Cronin et al., 2005; Hare et al., 2007; Martin et al., 2021; Melis et al., 2006), other-oriented prosocial behaviours (Burkart et al., 2014; Martin et al., 2021; Sehner et al., 2025), reproduction (Schülke & Ostner, 2008), and access to food resources (de Oliveira Terceiro et al., 2021; Fichtel et al., 2018; van Leeuwen et al., 2021). Social tolerance has also been shown to play an important role in social information transfer (Coussi-Korbel & Fragaszy, 1995; Dean et al., 2012; Whiten & van Schaik, 2007). Over recent decades, social tolerance has increasingly been linked to the facilitation of social learning across a variety of species, including chimpanzees (Nodé-Langlois et al., 2025; Whiten & van Schaik, 2007), orangutans (Fröhlich & van Schaik, 2022; Whiten & van Schaik, 2007), macaques (Garcia et al., 2022; Joly et al., 2017), capuchin monkeys (Coelho et al., 2024), lemurs (Sehner & Fichtel in prep.) and birds (Miller et al., 2014). Tolerating naïve individuals in close proximity becomes especially important when complex or difficult-to-acquire skills are transmitted, such as manufacturing and using tools (van Schaik et al., 1999) or abstract competencies like gestural communication (Fröhlich & van Schaik, 2022).

In this context, social tolerance is increasingly viewed as a potential driver of the development of single traditions and thus of cultural evolution (Burkart et al., 2009; van Schaik et al., 2019; Whiten & van Schaik, 2007). For instance, chimpanzees (*Pan spp*.) and orangutans (*Pongo spp*.) that spend more time in association show a more diverse cultural repertoire (Whiten & van Schaik, 2007). These results have recently been confirmed by Nodé-Langlois and colleagues (2025), who demonstrated that young chimpanzees benefit from living in more tolerant societies as they provide the opportunity to learn from multiple individuals, which in turn facilitates a rich repertoire of cultural behaviours. Furthermore, the development of new statistical tools, such as network-based diffusion analyses (NBDAs), has provided further insights into how behaviours spread within a society and showed that the likelihood of learning socially from conspecifics increases with physical proximity (Hoppitt et al., 2010).

In several species, the diffusion of innovations follow the structure of the social network: such as lobtail feeding in humpback whales (*Megaptera novaeangliae*) (Allen et al., 2013), the use of marine sponges as foraging tools in bottlenose dolphins (*Tursiops aduncus*) (Wild et al., 2019), the adoption of moss-sponging behaviour in chimpanzees (Hobaiter et al., 2014), the processing of novel food in vervet monkeys (*Chlorocebus pygerithrus*) (Canteloup et al., 2020), or the spread of an opening technique of feeding boxes in red-fronted lemurs (*Eulemur rufifrons*) and ring-tailed lemurs (*Lemur catta*) (Sehner & Fichtel, in prep). In addition, in squirrel monkeys (*Saimiri sciureus*), individuals with higher Eigenvector centrality (a measure of social connectedness) were more likely to acquire a novel foraging technique (Claidière et al., 2013). These findings highlight the importance of the structure of social ties within a network in shaping the transmission of socially learned behaviours.

A matter that has hitherto received less attention is how social tolerance may facilitate the initial emergence of an innovation. Innovations describe behavioural processes resulting in new or modified behaviours, including the ability to solve a novel problem (Reader & Laland, 2003). Social tolerance has been acknowledged to facilitate the emergence of material culture in primates (van Schaik et al., 1999) and it has been proposed that social tolerance can mediate the individual propensity to innovate (Sehner et al., 2022). Though individuals may intrinsically differ in their propensity to solve problems, leading to the idea that the probability that a novel problem is solved within a group is based on the available “pool of competences” or “skill pool” (Giraldeau, 1984; Morand-Ferron & Quinn, 2011). Hence, the more individuals a group contains, the more likely it is that a problem will be solved by one of these individuals, which additionally will be reinforced by mechanisms of social learning (Liker & Bókony, 2009; Morand-Ferron & Quinn, 2011; but see Cantor et al., 2020).

However, the collective problem-solving continuum hypothesis predicts that the likelihood of an individual solving a problem is not necessarily better in a group compared to when it is alone (Sehner et al., 2022). Based on the social tolerance of a species, the probability of solving a problem can either increase or decrease in a group setting. Groups with higher social tolerance are expected to be more prone to work cooperatively or at least not competitively on a problem, while individuals in less tolerant species will compete over access to a potential resource and block each other from focusing on the actual problem. Among other factors, this notion might explain why humans, which are regarded as hyper-cooperative (Burkart et al., 2014), have more complex innovations and thus a more complex cultural repertoire in comparison to other apes (Migliano & Vinicius, 2022; Sehner & Burkart, 2023). In nonhuman primates, it has been shown that highly socially tolerant primates like callitrichids excel in cooperative problem-solving (Cronin et al., 2005), and that the innovative complexity of groups surpasses that of individuals working alone (Sehner et al., 2022). In contrast, a study on chimpanzees has shown that the maximum complexity that individuals reach within a group reflects the maximum complexity that a single individual can achieve (Vale et al., 2021).

A key question in this context is whether all measures of social tolerance truly reflect the same underlying construct (DeTroy et al., 2022). In their review, these authors proposed a distinction between ‘behavioural social tolerance’, which refers to single tolerant (or intolerant) behaviour, and ‘structural social tolerance’, a multi-dimensional construct encompassing both affiliative and agonistic interactions that together reflect the overall hierarchical structure. Importantly, they argued that these constructs do not necessarily align within a species. This distinction also suggests that individual behaviours used to assess ‘behavioural social tolerance’ may not consistently correlate with each other, and that proximity patterns can differ substantially across social contexts. Nevertheless, it is common practice to use measures of tolerance taken outside the specific target context to assess social tolerance. For instance, social networks based on huddling or proximity have been used to test their role in facilitating social learning in foraging contexts (Coelho et al., 2024; Fröhlich & van Schaik, 2022; Garcia et al., 2022; Nodé-Langlois et al., 2025). Yet, assuming that tolerance is consistent across contexts can be misleading, as species like chimpanzees show tolerant behaviour when resources are difficult to monopolize (DeTroy et al., 2021; van Leeuwen et al., 2021, 2023), but exhibit a contrasting pattern when resources such as food puzzles can be monopolized (Knofe et al., 2019; Melis et al., 2011; Wittig & Boesch, 2003). Similar patterns have been observed in ring-tailed lemurs, where the monopolizability of food resources affects the proportion of group members permitted to co-feed (White et al., 2007).

To test whether tolerance is consistent across different foraging contexts and how it influences innovative behaviour, we conducted two complementary experiments. First, we measured social tolerance in a co-feeding paradigm (see for instance de Oliveira Terceiro et al., 2021; DeTroy et al., 2021; Fichtel et al., 2018), where food could not easily be monopolized by one individual. Second, we assessed tolerance during an extractive foraging task, using food puzzles that allowed dominant individuals to exclude others. We also quantified individual problem-solving propensity and examined whether, and in which contexts, social tolerance predicted successful food extractions. We tested white-faced capuchins (*Cebus capucinus*) and brown capuchins (*Sapajus apella*) to examine the relationship between social tolerance and innovation. While both species exhibit a diverse array of traditions (Fragaszy et al., 2004; Fragaszy & Perry, 2003), these traditions differ between these two taxa. Gracile capuchins (*Cebus spp*.) have a rich repertoire of social traditions (Perry, 2011, 2020; Perry et al., 2017), including behaviours like eye-poking where individuals show high tolerance toward each other. Although white-faced capuchins are less known for using tools (but see Barrett et al., 2018), they regularly engage in object use (Boinski, 1988). Robust capuchins (*Sapajus spp*.), on the other hand, have fewer documented social traditions but exhibit a wide range of foraging traditions, including tool-use (Falótico et al., 2022, 2024; Fragaszy et al., 2004; Valença et al., 2024) and show notable social tolerance in these contexts, tolerating close proximity of conspecifics. Together, these patterns raise two critical, yet underexplored issues. First, the observed differences in their repertoire of behavioural traditions may reflect a research bias: studies on gracile capuchins focusing more on social traditions, while those on robust capuchins emphasize foraging traditions, highlighting the need for more comparative research across these two taxa. Second, whether social tolerance in different contexts not only supports behavioural spread but also modulates its innovation remains unclear.

The aim of this study was to investigate the relationship between social tolerance in different contexts and innovative behaviour in two species of capuchins. We had no *a priori* prediction for potential differences in social tolerance between these two species. We expected that social tolerance in the co-feeding paradigm would be higher than in the extractive foraging paradigm, where resources can be more easily monopolized. However, we expected that on a dyadic level, tolerance between the two paradigms would correlate with each other, providing similar network structures within a group. Given that brown capuchins exhibit more foraging traditions than white-faced capuchins, we predicted that a larger number of brown capuchins would solve the test problems and that they would be overall more likely to solve more complex problems than white-faced capuchins. Lastly, we predicted a positive correlation between social tolerance and innovative propensity on an individual and a group level. We expected that more tolerant individuals would be more likely to solve the problem and that more tolerant groups contain more problem solvers. We expected this correlation to be stronger between social tolerance during extractive foraging and innovativeness than between social tolerance during co-feeding and innovativeness.

## Methods

### Study site and subjects

This study was conducted in white-faced and brown capuchins housed at the Primate Center of the University of Strasbourg, France. Experiments were conducted between October 2023 and January 2024. In total, we tested three groups of white-faced capuchins (n = 23 individuals in total) and three groups of brown capuchins (n = 20 individuals in total). Animals were housed either in outdoor enclosures overgrown with trees and shrubs or in outdoor enclosures enriched with branches, ropes, and diverse climbing structures. Both enclosure types had 24-hour access to indoor facilities that were also enriched with diverse climbing structures. Animals living in outdoor enclosures were housed indoors during winter. Animals had *ad libitum* access to water and were fed twice per day with commercial primate pellets. In addition, once per week, all groups were provided with fresh fruits and fresh vegetables. Data for three of the six groups (two brown- and one white-faced capuchin group) were collected between October and the beginning of November during a time where they still had access to the outdoor enclosures, whereas for the remaining three groups data were collected between December and January inside the indoor enclosures.

### Ethical statement

This non-invasive research complied with French legal requirements for the use of animals in research and EU Directive 2010/63/EU on the welfare of animals used for scientific purposes. The Strasbourg Ethics Committee for Animal Experiments (CREMEAS) confirmed that this study was ‘out of cope’ because of the absence of pain, constraint or nuisance for the animals as defined by Directive 2010/63 and included no ethical breach. Subjects participated voluntarily and were not deprived of food or water. The test area was not separated from the enclosure, which allowed the subjects to leave or join the area at will.

### Co-feeding experiment

To measure social tolerance, we used a co-feeding paradigm that has been used in other primate studies before (e.g., de Oliveira Terceiro et al., 2021; DeTroy et al., 2021; Fichtel et al., 2018). We prepared an area of 1 m^2^ (henceforth “co-feeding area”) per five animals in the group in which we distributed apple pieces. To mark the edges of the co-feeding area, we used naturally occurring stones of the enclosure or parts of the wooden enrichment within the enclosures. We used the same amount of apple pieces per group as for the procedure using the food puzzles (1 apple per 5 individuals). We chopped the apple into small cubes so that animals would not take larger pieces into their hands and leave the co-feeding area. We based this decision on our observations from the weekly fruit feeding, where animals took larger pieces and ran out of sight. In our approach, animals had to pick small food pieces from the ground, ensuring that feeding required them to remain inside the co-feeding area. A session started once the experimenter distributed the apple within the co-feeding area and ended when no individual was entering the area for five minutes or after a maximum of 30 minutes. We used the five minutes of no individual entering as a proximate estimation, indicating that all the food inside the co-feeding area was consumed. We recorded each session with two video cameras positioned outside the co-feeding area.

### Extractive foraging devices and procedure

We designed three versions of novel extractive foraging devices (henceforth food puzzles or puzzles) requiring one to three steps to acquire the food reward (**Fig. 1a-c**). For the first puzzle, animals had to rotate a door to either the left or right to access a hidden reward (1/24 of an apple) by reaching into a box (**Fig. 1a**). For the second puzzle, animals had to pull on a chain reaching out of a box, which moved a blockade out of the way so that they could push in a door and reach into the box (**Fig. 1b**). For the third puzzle, animals had to pull a metal rod blocking a slider that had to be pulled upwards and held in position to reach into the box and then pull on a chain to access the hidden reward (**Fig. 1c**).

**Figure 1.**
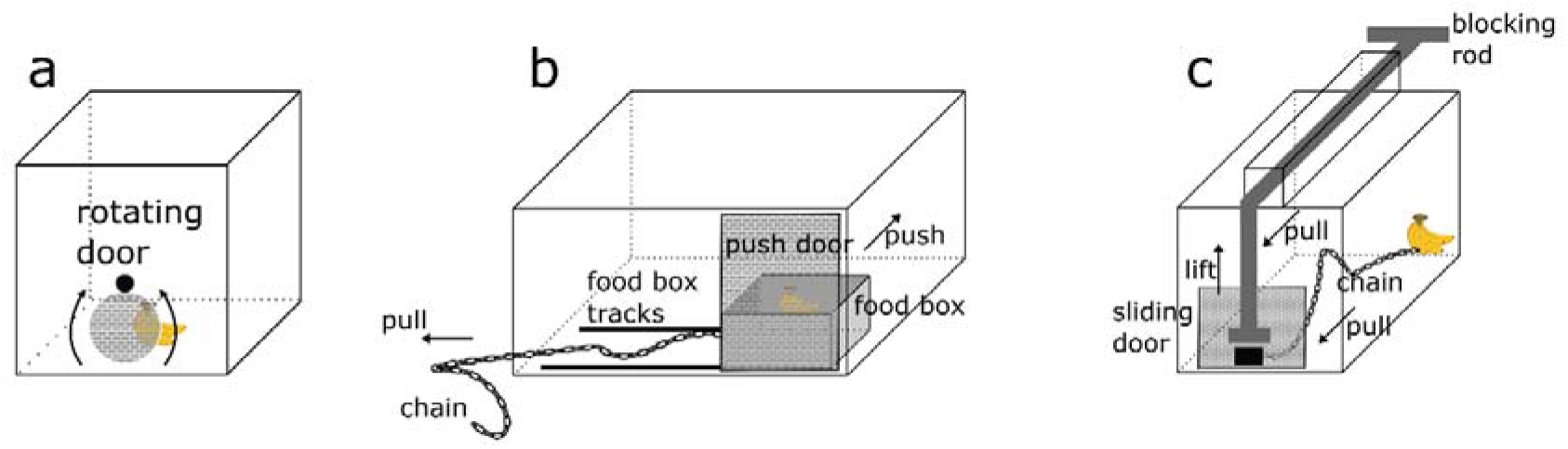
Schematic diagrams of the three extractive foraging devices. (a) Illustration of the one-step food puzzle, in which a rotating door had to be moved clockwise or counter-clockwise. (b) Illustration of the two-step food puzzle, in which a chain had to be pulled so that the food box moved to the side to unblock the push door; subsequently, animals had to push in the door to reach the food box. (c) Illustration of the three-step food puzzle, where animals had to push a metal rod to the side to unblock a sliding door. They then had to lift the door to reach and pull a chain, bringing the food reward within grabbing distance.

We presented the food puzzles always in the same order to ensure that if there were carry-over effects of learning from one puzzle to the next, it would be the same for all groups. Each puzzle was presented for five sessions on five consecutive days, and each session lasted 24 minutes, totalling 120 minutes of exposure per group per puzzle. Food puzzles were presented either in the morning or afternoon before the pellet feeding to ensure that individuals were motivated to participate. For the same reason, we tested animals before the weekly fruit feeding on the respective days. Before testing, the experimenter gave an audio cue that was associated with food to receive the attention of the individuals within the park. The puzzles were placed at the edges of the enclosure so that the experimenter could take place behind them to refill the food puzzles that were emptied from the other side of the fence. We prepared one apple per five adult individuals within a group by cutting it into 24 equal-sized pieces. A session started when a piece of apple was placed inside the puzzle and ended either after the 24 minutes were over or all food rewards were consumed. We also provided two replicas of each respective puzzle per session for the two larger groups (one group of white-faced and one group of brown capuchins) to reduce monopolization by a single individual. Although more puzzles would have been desirable, this was not possible due to logistical constraints, one of which was that the experimenter had to provide a new food item each time a puzzle was emptied. We recorded all sessions using two GoPro Hero9 black, which were placed outside of the enclosure.

### Neophobia

We also measured neophobia in an independent experiment using three novel objects. We used a red rubber ball with a small opening where this opening could only be explored with a single finger, a larger purple plastic ball with a large opening that allowed individuals to insert their hands to explore the item, and a teddy-like plush toy (**Fig. S1** in the Supplementary Material). We used three objects to test for repeatability of neophobia. Each group was presented with each object once for 20 minutes. We pseudorandomized the order in which the objects were presented. We recorded each session with a camera outside of the enclosure. Objects were fixed with zip ties to the enclosure in a way that animals could explore the objects and interact with them within a certain radius. However, objects could not be removed, which served the purpose that interactions could always be recorded and thus ensuring individual identification.

### Data analyses

All videos were analysed with Mangold interact Version 20.10.3.0 (**Table S1** in the Supplementary Material). Animals could be individually identified by the experimenter in front of the enclosure and the experimenter recorded the ID of each animal by naming them. From the video footage of the co-feeding experiment, we extracted the total time of an animal within the co-feeding area. For the extractive foraging devices, we recorded if, when, and how often an animal would approach the box (at least 1m proximity to a box; henceforth food puzzle area) and how long it would stay in the food puzzle area. We also recorded if, when, and how often an animal would explore the box and the duration of each exploration event. Lastly, we recorded if, when, and how often an animal would extract the hidden food reward. The video footage of the neophobia experiment was analysed in the same way, and we recorded which animal approached the novel object and when and for how long.

In the same way, we recorded if, when, and for how long an animal would interact with the novel object. However, interactions were overall rather rare. Hence, we used the approaching latency as a general measure of neophobia.

### Statistical analyses

Statistical analyses were carried out in R version 4.4.2 (R Core Team, 2024). We used the package “lme4” (Bates et al., 2014) for the generalized mixed models and the package “coxme” for the mixed effects Cox regression models (Therneau, 2024b). Figures were created either with the package “ggplot2” (Wickham, 2016) or the packages “survival” (Therneau, 2024a) and “survminer” (Kassambara et al., 2024). Social networks were created with the package “igraph” (Csárdi & Nepusz, 2006) and we used “vegan” (Oksanen et al., 2024) to compare network similarities. We used a likelihood ratio test for all GLMMs to test whether they predict the data better than a reduced model, including only random effects.

### Social tolerance

We fitted a generalized linear mixed model (GLMM) with a binomial error distribution to examine the likelihood of individuals being present in the experimental area (Model 1). The response was modelled as a two-column matrix of individuals being present and individuals being absent during each scan (every 15 seconds). Fixed effects included *species, paradigm* (co-feeding or extractive foraging), and the *z-transformed scan number*. Random effects included random intercepts and slopes for *condition* and *z-transformed scan number* within *group identity*, as well as random intercepts and slopes for *z-transformed scan number* within *session number* nested in *group identity*. This model allowed us to account for both temporal dynamics within sessions and repeated measures across groups and conditions.

We calculated a simple ratio association (SRA) score for each possible dyad within each group to create the social networks for each experimental condition (see van Leeuwen et al., 2021). The SRAs were calculated using the presence and absence of individuals within the co-feeding area and the presence and absence of individuals in the approaching area around the extractive foraging devices. We used the “fastgreedy” community detection algorithm to identify social networks within each group within each experimental treatment. We assessed the coefficient of variation for each group combining and each experimental treatment and performed a modified signed likelihood ratio test (MLSRT) to test for species differences. Comparing coefficients of variation can help reveal global tendencies of animals to be more selective than another group or in one condition compared to another. However, it cannot reveal whether networks are more similar to each other between conditions. It is possible that the overall coefficient remains rather stable while the connections between individuals have changed. To test this possibility, we used Mantel tests to compare networks within groups across conditions and assessed whether one species exhibited a stronger correlation than the other. Specifically, we compared SRA matrices using Pearson correlations and 10.000 permutations to determine statistical significance. We compared the resulting Mantel correlation coefficients between species using a two-sample t-test.

### Problem-solving

We analysed the approaching, exploring and solving behaviour separately. We calculated a series of mixed effects Cox regression models to analyse the latency of approaching, exploring and solving. We also calculated a series of GLMMs to analyse the approaching, exploration, and solving behaviour to assess differences between the two capuchin species. All models contained the same predictors and random effects, and analyses were conducted at the level of individual animals. Models included *species, puzzle, sex*, and *age* as predictors and *individual identity* nested in *group identity* as a random intercept. We specified a priori contrasts for our predictor *puzzle* to perform a polynomial trend analysis. We based these contrasts on our assumption that the food puzzles became increasingly difficult to solve from the first to the third. We first calculated the mixed effects Cox regression model to examine the latency of approaching (Model 2). We then calculated a GLMM with Poisson error distribution and log link function for the *number of approaching events* (Model 3) and a GLMM with a Gamma error Distribution and log link function to investigate the *average duration of an approach* (Model 4). Equally, we examined the latency of exploration (Model 5) the *number of exploration events* (Model 6) per individual and the *average duration of an exploration event* (Model 7). Lastly, we explored the latency of first time solving a puzzle (Model 8) and fitted a binomial GLMM with logit link function and the *number of extracted food items* and *the total number of possible items* as the response variable, to explore individual probabilities of solving the problem (Model 9).

We aimed to explore how social tolerance influences the probability of successfully extracting a food item in both species. However, the success-rate in white-faced capuchins was so low that we could perform the following analysis only within the brown capuchins. We calculated the Eigenvector centrality for each individual for each experimental treatment based on the respective SRA score. To assess whether neophobia toward objects reflects consistent individual differences, we examined behavioural repeatability as a proxy for personality. Due to limited consistency in neophobic responses across individual objects, we performed a principal component analysis (PCA), extracting a single score for the three neophobia tests, in which higher scores mean that individuals approached the novel objects on average later relative to those with lower scores. We fitted another binomial GLMM with logit-link function and the *number* of *extracted food items* and the *total number of possible items* as the response variables. This model included *age, sex*, and the two *Eigenvector centrality scores* (one for the co-feeding experiment and one for the food puzzle experiment) and the *first principal component for neophobia* as predictors (Model 10).

## Results

### Social tolerance

The full model (Model 1) explained the data better than the reduced model (likelihood ratio test: χ2 = 11.87; df = 3; *p* < 0.01; **Table S2** in the Supplementary Material). On average, the proportion of individuals of white-faced capuchins within the co-feeding area at each scan sample was around 8% and dropped to 5% during the food puzzle experiment. In brown capuchins, an average of 36% of group members were present in the co-feeding area, and 25% in the food puzzle are during each scan (**Fig. 2**). The model revealed that these differences between species and condition were significantly different from each other. Overall, significantly more group members of brown capuchins remained within the co-feeding and food puzzle areas compared to individuals of white-faced capuchins (odds ratio = 6.28, 95% CI = [1.99 – 19.85], *p* < 0.01). For both species, the condition mattered and the proportion of group members in the respective area was lower in the food puzzle condition compared to the co-feeding condition (odds ratio = 0.61, 95% CI = [0.44 – 0.84], *p* < 0.01). Lastly, time had a significant effect on the proportion of individuals within the respective area. Fewer individuals remained in the respective area the longer the experiment lasted (odds ratio = 0.78, 95% CI = [0.63 – 0.98], *p* = 0.03).

**Figure 2.**
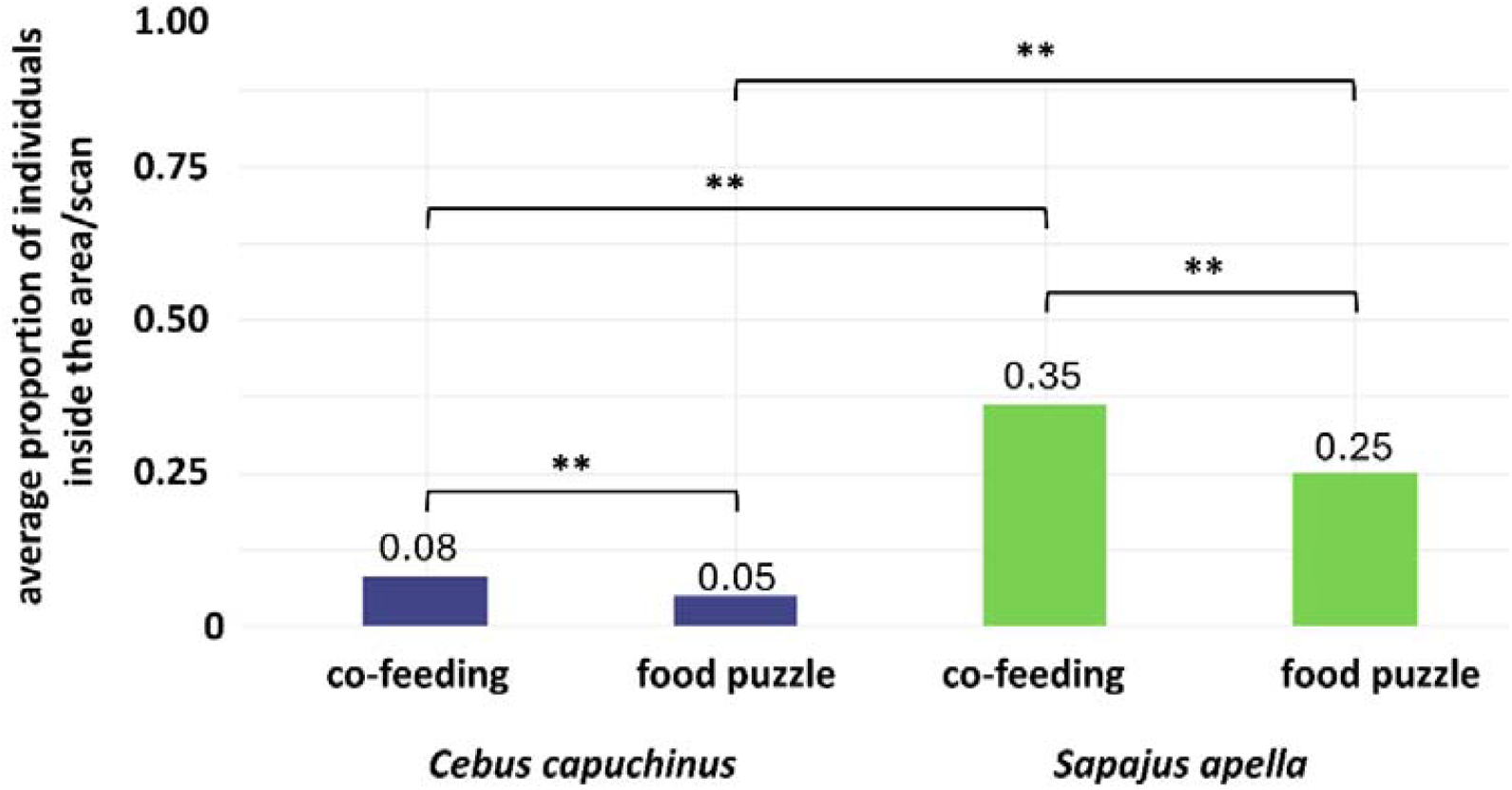
The proportion of individuals expected to be in the respective testing area (co-feeding area, or food puzzle area) at each scan interval. White-faced capuchins spent significantly less time in the areas than brown capuchins. In both species, tolerance was reduced in the food puzzle condition compared to the co-feeding condition.

In the next step, we calculated the social networks based on the simple ratio associations for both conditions (**Fig. S2 – S5** in the Supplementary Material). Comparing the coefficients of variation revealed that there was no difference between the two species (*C. capucinus*: CV = 2.34; *S. apella*: CV = 2.58; MSLRT = 0.19, *p* = 0.66). We also applied the MLSRT to test whether animals displayed greater selectivity in their social associations in the co-feeding condition compared to the extractive foraging condition. However, neither the white-faced (MSLRT = 1.76, *p* = 0.18) nor the brown capuchins (MSLRT = 0.22, *p* = 0.64) varied significantly between the two conditions. We also tested whether there was a significant difference in the coefficients of variation between conditions within species. Again, the coefficient of variation did not differ within the white-faced (MSLRT = 1.76, *p* = 0.18) nor within the brown capuchins (MSLRT = 0.22, *p* = 0.64).

Subsequently, we examined network similarities within groups, between conditions. The mean correlation between networks between brown (M = 0.89, SD =0.31) and white-faced capuchins (M = 0.07, SD =0.25) revealed a significant difference between the two species (Welch-test: t = −5.01, df =2.96, *p* = 0.01). Overall, the networks were more similar in the brown capuchins compared to the white-faced capuchins.

### Problem-solving

Overall, white-faced capuchins approached the food puzzles 869 times with an average approaching-duration of 22 seconds. During these approaches, they explored the puzzles 1859 times and successfully solved the puzzle 24 times out of 1800 opportunities. In contrast, brown capuchins approached the food puzzle area 705 times with an average duration of 148 seconds. Brown capuchins interacted 4397 times with the food puzzles and extracted a total of 1334 food items out of 1800 opportunities.

We first examined differences in approaching the food puzzle area (Models 2 – 4). There was no difference in the latency of approaching the food puzzle between species (**Fig. 3a; Table S3** in the Supplementary Material). However, older individuals were slightly faster approaching the food puzzle (proportional hazards ratio = 1.09, 95% CI = [1.03 – 1.14], *p* = 0.001). The full model (Model 3) explained the data better than the reduced model that included only random effects (likelihood ratio test: χ^2^ = 54.02, *p* < 0.001). The model indicates that *task complexity, sex*, and *age* significantly influenced the number of approaches (**Table S4** in the Supplementary Material). Individuals approached more complex tasks more often (odds ratio = 1.17, 95% CI = [1.07 – 1.28], *p* < 0.001). Furthermore, males approached the puzzles more often than females (odds ratio = 7.38, 95% CI = [2.65 – 20.54], *p* < 0.001) and older individuals more often than younger ones (odds ratio = 1.13, 95% CI = [1.05 – 1.23], *p* = 0.001). However, *species* had no effect on the number of approaches (*p* = 0.95). In contrast, a GLMM comparing the average time an individual stayed per visit revealed significance for age and species only (Model 4). Older individuals stayed longer per visit (odds ratio = 1.07, 95% CI = [1.03 – 1.11], *p* < 0.001) and brown capuchins stayed longer per visit than white-faced capuchins (odds ratio = 5.64, 95% CI = [3.39 – 9.36], *p* < 0.001; **Fig. 3b**).

**Figure 3.**
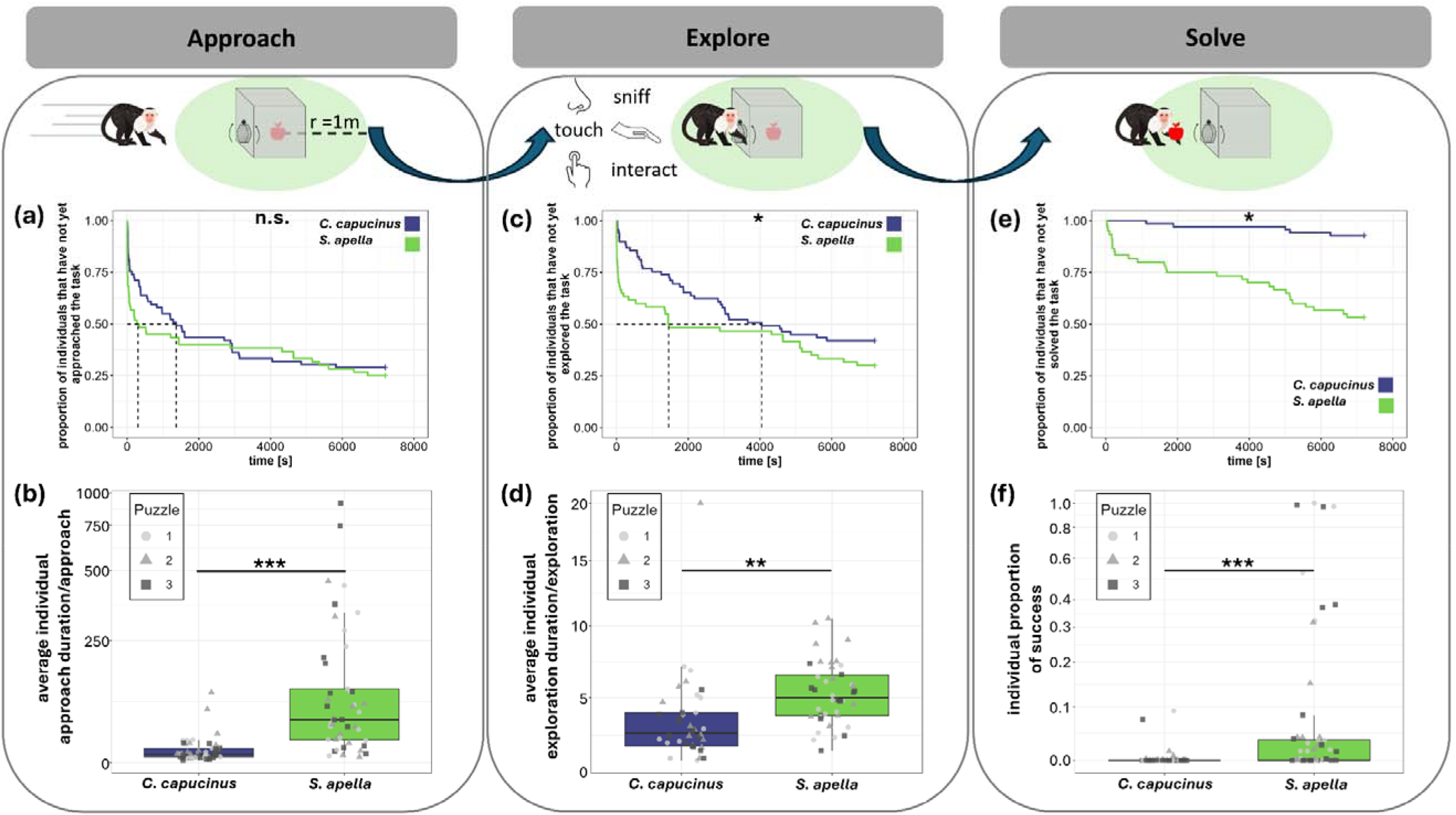
Overview of the extractive foraging task process in capuchin monkeys. The top section illustrates the 472 sequence of behaviours, from the initial approach to task exploration and eventual problem-solving. Panels (a–f) present quantitative measures of these behaviours: (a) latency to approach, (b) average approach duration per approach, (c) latency to explore, (d) average exploration duration per exploration event, (e) latency to solve, (f) individual success proportion based on total possible successes. Boxplots and survival curves are color-coded by species, with green representing *S. apella* and dark blue representing *C. capucinus*. In panels (b, d, f), puzzle identity is distinguished by shaped dots: circle (Puzzle 1), triangle (Puzzle 2), and square (Puz zle 3).

We then proceeded to examine the exploration behaviour (Models 5 – 7) In contrast to the approaching latency, we found a significant difference in exploration latency between the two species (**Fig. 3c; Table S5** in the Supplementary Material). Brown capuchins were on average faster than white-faced capuchins in exploring the food puzzles (proportional hazards ratio = 2.81, 95% CI = [1.08 – 7.28], *p* = 0.03). Also, older individuals explored the food puzzles faster (proportional hazards ratio = 1.17, 95% CI = [1.09 – 1.28], *p* < 0.001). Furthermore, we found an inverted U-shape relationship for approaching latency (proportional hazards ratio = 0.55, 95% CI = [0.35 – 0.87], *p* = 0.01). The full model examining the number of exploration events explained the data better than the reduced model (likelihood ratio test: χ^2^ = 89.91, *p* < 0.001; **Table S6** in the Supplementary Material). There was a trend for brown capuchins exploring more often than white-faced capuchins (*p* = 0.09). Equally to the number of approaching the food puzzles, older individuals (odds ratio = 1.28, 95% CI = [1.14 – 1.44], *p* < 0.001) and males (odds ratio = 18.11, 95% CI = [4.17 – 78.72], *p* < 0.001) explored food puzzles more often than younger ones and females. Our model revealed that the relationship between task complexity and explorative interactions was an inverted U-shape, suggesting that animals had fewer interactions during the first and third problems compared to the second problem (odds ratio = 0.84, 95% CI = [0.81 – 0.88], *p* < 0.001). Exploring the average duration of each exploration event revealed a similar result compared to the average duration of each approach (Supplementary S6). Brown capuchins, on average spent more time exploring the food puzzle per exploration event than white-faced capuchins (odds ratio = 1.81, 95% CI = [1.19 – 2.75], *p* < 0.01; **Fig. 3d**). Also, task complexity had an inverted U-shape effect on the average exploration duration per exploration event with longer exploration times during the second task compared to the other two (odds ratio = 0.76, 95% CI = [0.67 – 0.88], *p* < 0.001).

Lastly, we explored differences in solving the food puzzles (Models 8 – 9). Brown capuchins found the solution to a puzzle much faster than the white-faced capuchins did (proportional hazards ratio = 4.08, 95% CI = [1.24 – 13.39], *p* = 0.02; **Fig.3e**). In addition, older individuals were faster in extracting a food item from the puzzle than younger ones (proportional hazards ratio = 1.23, 95% CI = [1.12 – 1.34], *p* < 0.001; **Table S7** in the Supplementary Material). Subsequently, we analysed the proportion of successful extractions (**Table S8** in the Supplementary Material). The full model explained the data better than the reduced model (likelihood ratio test: χ^2^ = 3287, *p* < 0.001). Older individuals (odds ratio = 1.28, 95% CI = [1.14 – 1.44], *p* < 0.01) and males (odds ratio = 23.9, 95% CI = [1.4 – 407.2], *p* = 0.02) solved more problems than younger animals and females. Opposite to the pattern observed for explorative behaviour (inverted U-shape), problem-solving followed a U-shaped pattern, with more solutions found in the first and third tasks compared to the second task (odds ratio = 5.52, 95% CI = [4.67 – 6.55], *p* < 0.001). Brown capuchins were much more likely to solve a problem and extract a food reward than the white-faced capuchins (odds ratio = 555.8, 95% CI = [29.7 – 10404.9], *p* < 0.001; **Fig. 3f**).

### Social tolerance and innovativeness

Finally, we examined the relationship between social tolerance and problem-solving while also controlling for general neophobia (Model 10). Unfortunately, due to the small number of successful white-faced capuchins, we were only able to investigate which factors may have influenced individual solving success in brown capuchins. The repeatability analysis, conducted to assess individual consistency in neophobia across the three objects, showed that only 14% of the variation was explained by individual differences. Overall, neophobia did not differ between species (Mann-Whitney U test: W = 257, *p* = 0.52). The full model explained the data better than the null model (likelihood ratio test: χ^2^ = 14.36, *p* = 0.01). however, neither neophobia, nor the eigenvector centrality scores explained problem-solving success in the brown capuchins (**Table S9** in the Supplementary Material). Sex had also no influence on problem-solving success. In fact, only *age* revealed to have a significant influence on problem-solving success, with older individuals being more successful than younger individuals (odds ratio = 1.48, 95% CI = [1.21 – 1.81], *p* < 0.001).

## Discussion

### Social tolerance

Social tolerance has been shown to facilitate social learning in a variety of species. Here we measured social tolerance across contexts and tested its effect on innovativeness in two species of capuchin monkeys. We found that across contexts, brown capuchins were more tolerant than white-faced capuchins. In addition, social tolerance was more stable across contexts, and social networks showed a higher similarity in the brown than in the white faced-capuchins. We also found behavioural differences between the two species when approaching, exploring, and solving a novel food puzzle. Both species approached and explored the provided puzzles, but brown capuchins showed a higher persistence than white-faced capuchins. Overall, brown capuchins were much more successful problem-solvers. However, our proxies for social tolerance could not explain individual differences in solving success.

Brown capuchins showed a higher social tolerance than white-faced capuchins across contexts. We had no *a priori* prediction for species differences in social tolerance. Some studies reported that brown capuchins are highly tolerant (de Waal et al., 2008) and are almost as tolerant as common marmosets (Burkart & van Schaik, 2013), though less is known about tolerance levels in white-faced capuchins. However, given that white-faced capuchins engage in a variety of social traditions that require close proximity; for instance, eye-, nose-poking (Perry, 2011), this result is surprising. White-faced capuchins have been reported to be extremely tolerant towards each other during feeding contexts (Panger et al., 2002) and to allow other individuals to peer in close proximity, sniff, and even touch the food of their conspecifics (Perry & Rose, 1994) One possible explanation for this difference might be that white-faced capuchins are generally more arboreal than brown capuchins (Fragaszy et al., 2004). Since we conducted our social tolerance experiments on the ground, white-faced capuchins may have been more reluctant to participate in this experiment due to their more pronounced arboreal life-style. However, white-faced capuchins have been reported to be exceptionally terrestrial in the absence of predation pressure (Melis et al., 2014), and all individuals in our study population were observed regularly on the ground. Therefore, arboreality offers an insufficient explanation on its own.

Apart from species differences, social tolerance is also influenced by group composition (de Oliveira Terceiro et al., 2021; Kaufhold & van Leeuwen, 2019; van Leeuwen et al., 2021, 2023) and group size (Berghänel et al., 2025; Fichtel et al., 2018). However, since all groups of white-faced capuchins showed lower social tolerance than the groups of brown capuchins, and group size and composition are relatively similar, it is unlikely that this alone can explain the difference. For now, we have no definitive explanation of why brown capuchins were more socially tolerant. The lack of explanation is partly compounded by the fact that previous studies have emphasized different aspects of social behaviour in each species, limiting direct comparison. In particular, long-term observational work on white-faced capuchins has focused on social traditions (e.g., Perry, 2011; Perry et al., 2017), while studies on brown capuchins have often relied on experimental approaches and lack similarly detailed social data (e.g., Burkart & van Schaik, 2013, 2013; Custance et al., 1999). This observation highlights the need for comparative studies using consistent methodologies across species. As predicted, in both species social tolerance was higher in the co-feeding paradigm compared to the food puzzle paradigm. This difference is most likely due to the possibility of dominant individuals monopolizing the food puzzles, while it was more difficult for them to monopolize all pieces of the apple during the co-feeding experiment (see White et al., 2007 for similar results). This conclusion is supported by the fact that, across groups, mostly the dominant individuals approached and explored the food puzzles, while subordinate individuals approached only when the dominant animals were absent.

Since both measurements were taken in a foraging context and represent a specific tolerant behaviour (DeTroy et al., 2022), we predicted that levels of social tolerance would be correlated across conditions. Consistent with this notion, the coefficient of variation did not differ within species between conditions, suggesting that overall levels of social tolerance remained stable. However, a more detailed analysis of the social network revealed that, although individuals spent a similar amount of time interacting with others, the specific dyadic relationships had changed between conditions. The pattern recorded in this study matches observations in other robust capuchin species (*S. apella nigritus*), where individuals would groom the same individuals that they would spend time with during co-feeding (Tiddi et al., 2011). White-faced capuchins, on the other hand, have a less steep hierarchy, and patterns of association can be mixed across contexts and mostly depend on the sexes within a dyad (Crofoot et al., 2011; S. Perry, 1996, 1997, 1998). Importantly, it supports the idea of DeTroy and colleagues (2022) that measures of social tolerance within a population do not necessarily correlate with each other across contexts. Thus, whether social networks are stable across contexts seems to be influenced by species and maybe even populations. Comparable results have been reported in Barbary macaques, where huddling, but not grooming or proximity networks, predicted social learning in an extractive foraging paradigm (Garcia et al., 2022). It remains to be tested whether other forms of tolerance (e.g., proximity networks) might show stronger correlations with either the co-feeding network or the food puzzle network in the white-faced capuchin. Furthermore, it is unclear whether similar patterns would emerge in brown capuchins.

### Problem-solving

Both species approached the food puzzles an equal number of times, indicating that both species were at least interested in the puzzles. Moreover, the average latency to approach the task was similar between the two species, suggesting that species had an equally low level of neophobia. This result is supported by our independent neophobia test, where we also failed to find species differences. However, visits of white-faced capuchins in the approaching area of the food puzzles were much shorter than those of brown capuchins. One explanation may be that brown capuchins are more used to forage on the ground than white-faced capuchins are, and are thus more relaxed. However, in the enclosures animals are usually fed close to the ground. An alternative explanation is that brown capuchins tolerate each other more around the food puzzles, making them stay longer and allowing more individuals to stay simultaneously. In combination with a higher social tolerance, simple mechanisms like social enhancement can facilitate individuals of a group to spend more time around a novel object and ultimately facilitate perseverance (Fragaszy & Visalberghi, 1989; Sehner et al., 2022).

We found a similar pattern when examining the number of exploration events per species. Although there was a trend for brown capuchins to explore the food puzzles more often compared to the white-faced capuchins, a larger and also significant difference was found when examining the duration of the average exploration event. Brown capuchins explored much longer and started exploring earlier than the white-faced capuchins. A likely explanation is that brown capuchins were more motivated to interact with the devices, which could be triggered by simple social learning mechanisms like social facilitation (sensu Zajonc, 1965). The proportion of animals around the food puzzles was higher in brown capuchins, which may have triggered them to be more active and thus show more explorative behaviours, and for longer. The idea of socially induced perseverance also predicts that the success of others makes individuals more likely to approach and explore a food puzzle again (Sehner et al., 2022). However, in our study, the highest number of explorative events and the longest exploration duration per event were found during the second task, which showed the lowest number of solvers and was proportionally the puzzle with the least extracted food items. Again, simple social learning mechanisms may explain the motivation to explore the food puzzle, and over time, successful individuals required fewer actions to solve a puzzle for the first and third tasks, which decreased the overall number of explorative events and also the duration.

The largest difference between the two species was observed when it came to solving success. Brown capuchins were much more likely to successfully extract a food item from a puzzle. Due to their natural foraging behaviour and their propensity to use tools during extractive foraging (Fragaszy et al., 2004), brown capuchins were expected to be more successful than white-faced capuchins. However, we did not predict such a substantial difference between the two species. The required physiological and social traits to successfully solve novel problems are present in both species, since both are known to be rather tolerant during foraging and allow others to peer at their behaviour (Oppenheimer, 1968; Perry & Ordonez Jiménez, 2006). In addition, both species are described as highly explorative (Fragaszy et al., 2004; Perry et al., 2017) and have the required motor skills (Christel & Fragaszy, 2000; Torigoe, 1985). The latter is supported by the fact that at least some white-faced individuals solved the problems, of which one had missing digits on both hands due to injuries in the past. Moreover, although the differences in approaching and exploration duration may contribute to the result, they alone cannot explain the observed pattern. The average total exploration time of white-faced capuchins exceeded the average time brown capuchins spent exploring before solving the task for the first time. A possible explanation is that due to an overall higher tolerance in brown capuchins, the average individual could be more focused on the problem at hand and spend less time being socially vigilant, which according to the collective problem-solving continuum can enhance the average maximum achievable individual innovation complexity.

At the within-species level, however, we could only test brown capuchins on the influence of social tolerance on problem-solving abilities. Eigenvector centrality as a measure of how well an individual is connected socially to other individuals, also taking their connections into account, did not predict individual solving success. Neither the connectivity within the co-feeding network nor during the actual problem-solving sessions could explain individual success. This result is surprising and contrasts with previous studies showing that more tolerant capuchins were more likely to solve a novel problem (Morton et al., 2021). The values for Eigenvector centrality were based on the simple ratio associations given through the time individuals spend in the area around the foraging devices or within the co-feeding area. However, single individuals monopolized the foraging devices for a large proportion of the sessions, meaning that individuals may be well connected within the network and tolerated around the food sources, but still had little access to the actual device. In more natural settings, where resources are usually more scattered, individuals may benefit from tolerant groups and learn how to access certain resources. However, due to our setup, individuals may have learned to solve the problem, but could not access it due to more dominant individuals monopolizing the apparatuses. Overall, our results fit with the general assumption that social enhancement facilitates innovation in capuchins, while restricted access inhibits innovation (Fragaszy & Visalberghi, 1989). Additionally, we cannot rule out that our small sample size masks the effect of social tolerance on individual innovativeness. To disentangle these effects, future studies should test single individuals of different ranks as a control condition and also compare equal numbers of single individuals with groups, in order to separate rank-related effects from the numerical advantages of larger groups (Cantor et al., 2020; Sehner et al., 2022).

Ultimately, the best predictor for individual problem solving success was age, with older individuals being more successful compared to younger ones. These results are in contrast to earlier observations both in the wild and captivity, where younger individuals tended to invent more novel behaviours regarding foraging than older ones (Morton et al., 2021; Perry et al., 2017). However, captive individuals often behave differently from their wild counterparts and show differences between sites due to several possible captivity effects (Clubb & Mason, 2003; de Oliveira Terceiro et al., 2021; Fichtel et al., 2018; Splinter et al., 2025). For once, the cognitive load in captive animals might be reduced due to a lack of predation pressure or the necessity to forage for food, and free resources to explore novel items (Forss et al., 2015). However, this should be the case for both, older and younger individuals. Another explanation is that animals in research facilities are tested repeatedly and older individuals have more experience in these scenarios and expect a reward for participating, whereas younger individuals have less experience and are also then blocked by their older conspecifics. This notion is supported by the fact that older individuals also had longer approaches, longer exploration durations, and shorter latencies to explore than younger individuals. Ultimately, the successful older individuals in our experiment were also the most dominant individuals across conditions, which explains why males were more innovative than females.

Lastly, we cannot rule out the possibility that innovations happened only once in each group and were subsequently spread via social learning. Given their strong tendency to learn from one another (Coelho et al., 2015; Custance et al., 1999; Kuroshima et al., 2008), at least some solvers probably acquired the solution socially rather than independently. However, there were two stable subgroups in the largest group of brown capuchins, which did not explore the food puzzles at the same time, making social transmission for one of the puzzles improbable, and even and even less so across all three tasks. Hence, at least some individuals invented the solution independently. A network-based diffusion analysis could resolve this question and help explain to what extent social learning is involved. Unfortunately, due to the small sample size, with only three groups of brown capuchins available and two of them being small, it was not feasible to conduct a meaningful Order of Acquisition Analysis (Franz & Nunn, 2010). Nevertheless, the pronounced difference between white-faced and brown capuchins in both problem-solving success and social tolerance suggests that social tolerance modulates not only the likelihood of social learning but also the propensity for individual innovation.

In conclusion, our study adds to the body of research suggesting that social tolerance can facilitate both, innovativeness and social learning and, hence, cultural evolution. Our study also contributes to our understanding of why brown capuchins show a higher repertoire of foraging traditions than white-faced capuchins since they show higher social tolerance in feeding and foraging contexts. It raises the question of whether white-faced capuchins are more tolerant in other contexts which facilitates the emergence of social traditions. Future work should adopt the approach of DeTroy and colleagues (2022) to measure tolerance as a structural concept in white-faced capuchins and to explore differences with other species. Furthermore, future work should also test whether these capuchin species are more or less innovative within a group compared to when they are alone.

As suggested by the “collective problem-solving continuum”, brown capuchins are expected to show a higher propensity to innovate in a group setting, whereas white-faced capuchins may work better alone without the interference of conspecifics (Sehner et al., 2022). By advancing our understanding of how social tolerance shapes both innovation and information diffusion, this line of research contributes to our understanding of the social foundations of cultural evolution across taxa and ultimately, the evolutionary roots of cumulative culture in humans.

## Acknowledgements

We are grateful to Silabe and the Centre de Primatology Niederhausbergen for providing the facilities to conduct this research. We thank the animal caretakers and the animal welfare management at the Centre de Primatologie for helping with the handling of the animals and their advice. We are particular grateful to Gael Raimbault and Ruta Vaicekauskaite for their help. We also thank Alice Beaud for helping us to identify the animals at the beginning of the experiment. We are grateful to Roger Mundry for fruitful discussions about our statistical approach and helpful advice. CF thanks the Collaborative Research Center “Cognition of Interaction” - SFB 1528, funded by the German Research Foundation (DFG) for discussions. This study was funded by an SNF Postdoc.Mobility Grant (grant number P500PB_217864/1) to Sandro Sehner.

## References

Allen, J., Weinrich, M., Hoppitt, W., & Rendell, L. (2013). Network-based diffusion analysis reveals cultural transmission of lobtail feeding in humpback whales. Science, 340(6131), 485–488. 10.1126/science.1231976

Barrett, B. J., Monteza-Moreno, C. M., Dogandžić, T., Zwyns, N., Ibáñez, A., & Crofoot, M. C. (2018). Habitual stone-tool-aided extractive foraging in white-faced capuchins, Cebus capucinus. Royal Society Open Science, 5(8), 181002. 10.1098/rsos.181002

Bates, D., Mächler, M., Bolker, B., & Walker, S. (2014). Fitting Linear Mixed-Effects Models using lme4 (arXiv:1406.5823). arXiv. 10.48550/arXiv.1406.5823

Berghänel, A., Lazzaroni, M., Ferenc, M., Pilot, M., El Berbri, I., Marshall-Pescini, S., & Range, F. (2025). Cofeeding at rich clumped food patches in free-ranging dogs: Social tolerance or scramble competition? Behavioral Ecology and Sociobiology, 79(4). 10.1007/s00265-025-03590-8

Boinski, S. (1988). Use of a club by a wild white-faced capuchin (Cebus capucinus) to attack a venomous snake (Bothrops asper). American Journal of Primatology, 14(2), 177–179. 10.1002/ajp.1350140208

Burkart, J. M., Allon, O., Amici, F., Fichtel, C., Finkenwirth, C., Heschl, A., Huber, J., Isler, K., Kosonen, Z. K., Martins, E., Meulman, E. J., Richiger, R., Rueth, K., Spillmann, B., Wiesendanger, S., & van Schaik, C. P. (2014). The evolutionary origin of human hyper-cooperation. Nature Communications, 5(1), 4747. 10.1038/ncomms5747

Burkart, J. M., Hrdy, S. B., & Van Schaik, C. P. (2009). Cooperative breeding and human cognitive evolution. Evolutionary Anthropology: Issues, News, and Reviews, 18(5), 175–186. 10.1002/evan.20222

Burkart, J. M., & van Schaik, C. (2013). Group service in macaques (Macaca fuscata), capuchins (Cebus apella) and marmosets (Callithrix jacchus): A comparative approach to identifying proactive prosocial motivations. Journal of Comparative Psychology, 127(2), 212–225. 10.1037/a0026392

Canteloup, C., Hoppitt, W., & van de Waal, E. (2020). Wild primates copy higher-ranked individuals in a social transmission experiment. Nature Communications, 11(1), 459. 10.1038/s41467-019-14209-8

Cantor, M., Aplin, L. M., & Farine, D. R. (2020). A primer on the relationship between group size and group performance. Animal Behaviour, 166, 139–146. 10.1016/j.anbehav.2020.06.017

Christel, M. I., & Fragaszy, D. (2000). Manual function in Cebus apella. Digital mobility, preshaping, and endurance in repetitive grasping. International Journal of Primatology, 21(4), 697–719. 10.1023/A:1005521522418

Claidière, N., Messer, E. J. E., Hoppitt, W., & Whiten, A. (2013). Diffusion dynamics of socially learned foraging techniques in squirrel monkeys. Current Biology, 23(13), 1251–1255. 10.1016/j.cub.2013.05.036

Clubb, R., & Mason, G. (2003). Captivity effects on wide-ranging carnivores. Nature, 425(6957), 473–474. 10.1038/425473a

Coelho, C. G., Falótico, T., Izar, P., Mannu, M., Resende, B. D., Siqueira, J. O., & Ottoni, E. B. (2015). Social learning strategies for nut-cracking by tufted capuchin monkeys (Sapajus spp.). Animal Cognition, 18(4), 911–919. 10.1007/s10071-015-0861-5

Coelho, C. G., Garcia-Nisa, I., Ottoni, E. B., & Kendal, R. L. (2024). Social tolerance and success-biased social learning underlie the cultural transmission of an induced extractive foraging tradition in a wild tool-using primate. Proceedings of the National Academy of Sciences, 121(48), e2322884121. 10.1073/pnas.2322884121

Coussi-Korbel, S., & Fragaszy, D. M. (1995). On the relation between social dynamics and social learning. Animal Behaviour, 50(6), 1441–1453. 10.1016/0003-3472(95)80001-8

Crofoot, M. C., Rubenstein, D. I., Maiya, A. S., & Berger-Wolf, T. Y. (2011). Aggression, grooming and group-level cooperation in white-faced capuchins (Cebus capucinus): Insights from social networks. American Journal of Primatology, 73(8), 821–833. 10.1002/ajp.20959

Cronin, K. A., Kurian, A. V., & Snowdon, C. T. (2005). Cooperative problem solving in a cooperatively breeding primate (Saguinus oedipus). Animal Behaviour, 69(1), 133–142. 10.1016/j.anbehav.2004.02.024

Csárdi G. & Nepusz T. (2006). The igraph software package for complex network research. Igraph – the Network Analysis Package. https://igraph.org/

Custance, D., Whiten, A., & Fredman, T. (1999). Social learning of an artificial fruit task in capuchin monkeys (Cebus apella). Journal of Comparative Psychology, 113(1), 13–23. 10.1037/0735-7036.113.1.13

de Oliveira Terceiro, F. E., Arruda, M. de F., van Schaik, C. P., Araújo, A., & Burkart, J. M. (2021). Higher social tolerance in wild versus captive common marmosets: The role of interdependence. Scientific Reports, 11(1), 825. 10.1038/s41598-020-80632-3

De Waal, F. B. M. (1986). Class structure in a rhesus monkey group: The interplay between dominance and tolerance. Animal Behaviour, 34(4), 1033–1040. 10.1016/S0003-3472(86)80162-2

de Waal, F. B. M., Leimgruber, K., & Greenberg, A. R. (2008). Giving is self-rewarding for monkeys. Proceedings of the National Academy of Sciences, 105(36), 13685–13689. 10.1073/pnas.0807060105

Dean, L., Hoppitt, W., Rendell, L., & Webster, M. M. (2012). Social Learning Strategies, Mechanisms, and Models. In E. A. Wasserman & T. R. Zentall (Eds.), The Oxford Handbook of Comparative Cognition (pp. 832–850). Oxford University Press. 10.1093/oxfordhb/9780195392661.013.0042

DeTroy, S. E., Haun, D. B. M., & van Leeuwen, E. J. C. (2022). What isn’t social tolerance? The past, present, and possible future of an overused term in the field of primatology. Evolutionary Anthropology: Issues, News, and Reviews, 31(1), 30–44. 10.1002/evan.21923

DeTroy, S. E., Ross, C. T., Cronin, K. A., van Leeuwen, E. J. C., & Haun, D. B. M. (2021). Cofeeding tolerance in chimpanzees depends on group composition: A longitudinal study across four communities. iScience, 24(3), 102175. 10.1016/j.isci.2021.102175

Falótico, T., Macedo, A. C., de Jesus, M. A., Espinola, T., & Valença, T. (2024). Nutcracking success and efficiency in two wild capuchin monkey populations. Royal Society Open Science, 11(6), 240161. 10.1098/rsos.240161

Falótico, T., Valença, T., Verderane, M. P., & Fogaça, M. D. (2022). Stone tools differences across three capuchin monkey populations: Food’s physical properties, ecology, and culture. Scientific Reports, 12(1), 14365. 10.1038/s41598-022-18661-3

Fichtel, C., Schnoell, A. V., & Kappeler, P. M. (2018). Measuring social tolerance: An experimental approach in two lemurid primates. Ethology, 124(1), 65–73. 10.1111/eth.12706

Forss, S. I. F., Schuppli, C., Haiden, D., Zweifel, N., & van Schaik, C. P. (2015). Contrasting responses to novelty by wild and captive orangutans: Novelty Response in Orangutans. American Journal of Primatology, 77(10), 1109–1121. 10.1002/ajp.22445

Fragaszy, D. M., & Perry, S. (2003). Towards a biology of traditions. In D. M. Fragaszy & S. Perry (Eds.), The Biology of Traditions: Models and Evidence (pp. 1–32). Cambridge University Press. 10.1017/CBO9780511584022.002

Fragaszy, D. M., & Visalberghi, E. (1989). Social influences on the acquisition of tool-using behaviors in tufted capuchin monkeys (Cebus apella). Journal of Comparative Psychology, 103(2), 159–170. 10.1037/0735-7036.103.2.159

Fragaszy, D. M., Visalberghi, E., & Fedigan, L. M. (2004). The Complete Capuchin: The Biology of the Genus Cebus. Cambridge University Press.

Franz, M., & Nunn, C. L. (2010). Investigating the impact of observation errors on the statistical performance of network-based diffusion analysis. Learning & Behavior, 38(3), 235–242. 10.3758/LB.38.3.235

Fröhlich, M., & van Schaik, C. P. (2022). Social tolerance and interactional opportunities as drivers of gestural redoings in orang-utans. Philosophical Transactions of the Royal Society B: Biological Sciences, 377(1859), 20210106. 10.1098/rstb.2021.0106

Garcia, A. C., Parsons, M. A., & Young, J. K. (2022). Effects of early-life experience on innovation and problem-solving in captive coyotes. Behavioral Ecology and Sociobiology, 76(10), 141. 10.1007/s00265-022-03251-0

Giraldeau, L.-A. (1984). Group foraging: The skill pool effect and frequencydependent learning. The American Naturalist, 124(1), 72–79. 10.1086/284252

Hare, B., Melis, A. P., Woods, V., Hastings, S., & Wrangham, R. (2007). Tolerance allows bonobos to outperform chimpanzees on a cooperative task. Current Biology, 17(7), 619–623. 10.1016/j.cub.2007.02.040

Hobaiter, C., Poisot, T., Zuberbühler, K., Hoppitt, W., & Gruber, T. (2014). Social network analysis shows direct evidence for social transmission of tool use in wild chimpanzees. PLoS Biology, 12(9), e1001960. 10.1371/journal.pbio.1001960

Hoppitt, W., Boogert, N. J., & Laland, K. N. (2010). Detecting social transmission in networks. Journal of Theoretical Biology, 263(4), 544–555. 10.1016/j.jtbi.2010.01.004

Joly, M., Micheletta, J., De Marco, A., Langermans, J. A., Sterck, E. H. M., & Waller, B. M. (2017). Comparing physical and social cognitive skills in macaque species with different degrees of social tolerance. Proceedings of the Royal Society B: Biological Sciences, 284(1862), 20162738. 10.1098/rspb.2016.2738

Kassambara, Alboukadel, Kosinski, Marcin, & Biecek, Przemyslaw. (2024). survminer: Drawing Survival Curves using “ggplot2.” https://CRAN.R-project.org/package=survminer

Kaufhold, S. P., & van Leeuwen, E. J. C. (2019). Why intergroup variation matters for understanding behaviour. Biology Letters, 15(11), 20190695. 10.1098/rsbl.2019.0695

Knofe, H., Engelmann, J., Tomasello, M., & Herrmann, E. (2019). Chimpanzees monopolize and children take turns in a limited resource problem. Scientific Reports, 9(1), 7597. 10.1038/s41598-019-44096-4

Kuroshima, H., Kuwahata, H., & Fujita, K. (2008). Learning from others’ mistakes in capuchin monkeys (Cebus apella). Animal Cognition, 11(4), 599–609. 10.1007/s10071-008-0150-7

Liker, A., & Bókony, V. (2009). Larger groups are more successful in innovative problem solving in house sparrows. Proceedings of the National Academy of Sciences, 106(19), 7893–7898. 10.1073/pnas.0900042106

Martin, J. S., Koski, S. E., Bugnyar, T., Jaeggi, A. V., & Massen, J. J. M. (2021). Prosociality, social tolerance and partner choice facilitate mutually beneficial cooperation in common marmosets, Callithrix jacchus. Animal Behaviour, 173, 115–136. 10.1016/j.anbehav.2020.12.016

Melis, A. P., Hare, B., & Tomasello, M. (2006). Chimpanzees recruit the best collaborators. Science, 311(5765), 1297–1300. 10.1126/science.1123007

Melis, A. P., Schneider, A.-C., & Tomasello, M. (2011). Chimpanzees, Pan troglodytes, share food in the same way after collaborative and individual food acquisition. Animal Behaviour, 82(3), 485–493. 10.1016/j.anbehav.2011.05.024

Migliano, A. B., & Vinicius, L. (2022). The origins of human cumulative culture: From the foraging niche to collective intelligence. Philosophical Transactions of the Royal Society B: Biological Sciences, 377(1843), 20200317. 10.1098/rstb.2020.0317

Miller, R., Schiestl, M., Whiten, A., Schwab, C., & Bugnyar, T. (2014). Tolerance and social facilitation in the foraging behaviour of free-ranging crows (Corvus corone corone; C. c. Cornix). Ethology, 120(12), 1248–1255. 10.1111/eth.12298

Morand-Ferron, J., & Quinn, J. L. (2011). Larger groups of passerines are more efficient problem solvers in the wild. Proceedings of the National Academy of Sciences, 108(38), 15898–15903. 10.1073/pnas.1111560108

Morton, F. B., Buchanan-Smith, H. M., Brosnan, S. F., Thierry, B., Paukner, A., Essler, J. L., Marcum, C. S., & Lee, P. C. (2021). Studying animal innovation at the individual level: A ratings-based assessment in capuchin monkeys (Sapajus [Cebus] sp.). Journal of Comparative Psychology, 135(2), 258–265. 10.1037/com0000264

Nodé-Langlois, O., Rolland, E., Girard-Buttoz, C., Samuni, L., Ferrari, P. F., Wittig, R. M., & Crockford, C. (2025). Social tolerance and role model diversity increase tool use learning opportunities across chimpanzee ontogeny. Communications Biology, 8(1), 1–17. 10.1038/s42003-025-07885-4

Oksanen, J., Kindt, R., Legendre, P., O’Hara, B., Simpson, G., Solymos, P., Henry, M., Stevens, H., Maintainer, H., & Oksanen, M. J. (2024). The vegan Package. https://CRAN.R-project.org/package=vegan

Oppenheimer, J. R. (1968). Behavior and ecology of the white-faced monkey, Cebus capucinus, on Barro Colorado Island, Canal Zone. University of Illinois at Urbana-Champaign. https://www.proquest.com/openview/fe99f003fa97376939ce1be1251d6b36/1?cbl=18750&diss=y&pq-origsite=gscholar

Panger, M. A., Perry, S., Rose, L., Gros-Louis, J., Vogel, E., Mackinnon, K. C., & Baker, M. (2002). Cross-site differences in foraging behavior of white-faced capuchins (Cebus capucinus). American Journal of Physical Anthropology, 119(1), 52–66. 10.1002/ajpa.10103

Perry, S. (1996). Female-female social relationships in wild white-faced capuchin monkeys, Cebus capucinus. American Journal of Primatology, 40(2), 167–182. 10.1002/(SICI)1098-2345(1996)40:2<167::AID-AJP4>3.0.CO;2-W

Perry, S. (1997). Male-female social relationships in wild white-faced capuchins (Cebus capucinus). Behaviour, 134(7–8), 477–510. 10.1163/156853997X00494

Perry, S. (1998). Male-male social relationships in wild white-faced capuchins (Cebus capucinus). Behaviour, 135(2), 139–172.

Perry, S. (2011). Social traditions and social learning in capuchin monkeys (Cebus). Philosophical Transactions of the Royal Society B: Biological Sciences, 366(1567), 988–996. 10.1098/rstb.2010.0317

Perry, S. (2020). Behavioural variation and learning across the lifespan in wild white-faced capuchin monkeys. Philosophical Transactions of the Royal Society B: Biological Sciences, 375(1803), 20190494. 10.1098/rstb.2019.0494

Perry, S. E., Barrett, B. J., & Godoy, I. (2017). Older, sociable capuchins (Cebus capucinus) invent more social behaviors, but younger monkeys innovate more in other contexts. Proceedings of the National Academy of Sciences, 114(30), 7806–7813. 10.1073/pnas.1620739114

Perry, S., & Ordonez Jiménez, J. (2006). The effects of food size, rarity, and processing complexity on white-faced capuchins visual attention to foraging conspecifics. In R. W. Sussman & D. R. Phillips (Eds.), Feeding Ecology in Apes and other Primates (pp. 203–234). Univ. Press. https://pure.mpg.de/pubman/faces/ViewItemOverviewPage.jsp?itemId=item_1555062

Perry, S., & Rose, L. (1994). Begging and transfer of coati meat by white-faced capuchin monkeys,Cebus capucinus. Primates, 35(4), 409–415. 10.1007/BF02381950

R Core Team. (2024). R: A Language and Environment for Statistical Computing (Version 2024) [Computer software]. R Foundation for Statistical Computing. https://www.R-project.org/

Reader, S. M., & Laland, K. N. (2003). Animal Innovation. Oxford University Press.

Rina Evasoa, M., Zimmermann, E., Hasiniaina, A. F., Rasoloharijaona, S., Randrianambinina, B., & Radespiel, U. (2019). Sources of variation in social tolerance in mouse lemurs (Microcebus spp.). BMC Ecology, 19(1), 20. 10.1186/s12898-019-0236-x

Schülke, O., & Ostner, J. (2008). Male reproductive skew, paternal relatedness, and female social relationships. American Journal of Primatology, 70(7), 695–698. 10.1002/ajp.20546

Sehner, S., & Burkart, J. M. (2023). Cumulative culture. Zeitschrift Für Entwicklungspsychologie Und Pädagogische Psychologie, 55(1), 9–13. 10.1026/0049-8637/a000268

Sehner, S., Willems, E. P., Baumeyer, A., Davis, L., van Schaik, C. P., & Burkart, J. M. (2025). Sensitivity to immature skill deficits. Food sharing experiments in squirrel monkeys (Saimiri boliviensis) and common marmosets (Callithrix jacchus). Journal of Comparative Psychology, 139(3), 178–191. 10.1037/com0000399

Sehner, S., Willems, E. P., Vinicus, L., Migliano, A. B., van Schaik, C. P., & Burkart, J. M. (2022). Problem-solving in groups of common marmosets (Callithrix jacchus): More than the sum of its parts. PNAS Nexus, 1(4), pgac168. 10.1093/pnasnexus/pgac168

Splinter, F., Fichtel, C., & Radespiel, U. (2025). Cognitive performance in wild and captive grey mouse lemurs (Microcebus murinus). International Journal of Primatology, 46(3), 644–663. 10.1007/s10764-025-00490-6

Therneau, T. M. (2024a). A Package for Survival Analysis in S. R package version 3. 8-3, 83.

Therneau, T. M. (2024b). coxme: Mixed Effects Cox Models (Version R package version 2. 2-22). https://CRAN.R-project.org/package=coxme

Tiddi, B., Aureli, F., Polizzi di Sorrentino, E., Janson, C. H., & Schino, G. (2011). Grooming for tolerance? Two mechanisms of exchange in wild tufted capuchin monkeys. Behavioral Ecology, 22(3), 663–669. 10.1093/beheco/arr028

Torigoe, T. (1985). Comparison of object manipulation among 74 species of nonhuman primates. Primates, 26(2), 182–194. 10.1007/BF02382017

Vale, G. L., McGuigan, N., Burdett, E., Lambeth, S. P., Lucas, A., Rawlings, B., Schapiro, S. J., Watson, S. K., & Whiten, A. (2021). Why do chimpanzees have diverse behavioral repertoires yet lack more complex cultures? Invention and social information use in a cumulative task. Evolution and Human Behavior, 42(3), 247–258. 10.1016/j.evolhumbehav.2020.11.003

Valença, T., Oliveira Affonço, G., & Falótico, T. (2024). Wild capuchin monkeys use stones and sticks to access underground food. Scientific Reports, 14(1), 10415. 10.1038/s41598-024-61243-8

van de Waal, E., Renevey, N., Favre, C. M., & Bshary, R. (2010). Selective attention to philopatric models causes directed social learning in wild vervet monkeys. Proceedings of the Royal Society B: Biological Sciences, 277(1691), 2105–2111. 10.1098/rspb.2009.2260

van Leeuwen, E. J. C., Staes, N., Brooker, J. S., Kordon, S., Nolte, S., Clay, Z., Eens, M., & Stevens, J. M. G. (2023). Group-specific expressions of co-feeding tolerance in bonobos and chimpanzees preclude dichotomous species generalizations. iScience, 26(12), 108528. 10.1016/j.isci.2023.108528

van Leeuwen, E. J. C., Van Donink, S., Eens, M., & Stevens, J. M. G. (2021). Group-level variation in co-feeding tolerance between two sanctuary-housed communities of chimpanzees (Pan troglodytes). Ethology, 127(7), 517–526. 10.1111/eth.13154

van Schaik, C. P., Deaner, R. O., & Merrill, M. Y. (1999). The conditions for tool use in primates: Implications for the evolution of material culture. Journal of Human Evolution, 36(6), 719–741. 10.1006/jhev.1999.0304

van Schaik, C. P., Pradhan, G. R., & Tennie, C. (2019). Teaching and curiosity: Sequential drivers of cumulative cultural evolution in the hominin lineage. Behavioral Ecology and Sociobiology, 73(1), 2. 10.1007/s00265-018-2610-7

White, F. J., Overdorff, D. J., Keith-Lucas, T., Rasmussen, M. A., Eddie Kallam, W., & Forward, Z. (2007). Female dominance and feeding priority in a prosimian primate: Experimental manipulation of feeding competition. American Journal of Primatology, 69(3), 295–304. 10.1002/ajp.20346

Whiten, A., & van Schaik, C. P. (2007). The evolution of animal ‘cultures’ and social intelligence. Philosophical Transactions of the Royal Society B: Biological Sciences, 362(1480), 603–620. 10.1098/rstb.2006.1998

Wickham, H. (2016). ggplot2: Elegant graphics for Data Analysis. In H. Wickham (Ed.), Ggplot2: Elegant Graphics for Data Analysis (pp. 189–201). Springer International Publishing. 10.1007/978-3-319-24277-4_9

Wild, S., Allen, S. J., Krützen, M., King, S. L., Gerber, L., & Hoppitt, W. J. E. (2019). Multi-network-based diffusion analysis reveals vertical cultural transmission of sponge tool use within dolphin matrilines. Biology Letters, 15(7), 20190227. 10.1098/rsbl.2019.0227

Wittig, R. M., & Boesch, C. (2003). Food competition and linear dominance hierarchy among female chimpanzees of the Taï National Park. International Journal of Primatology, 24(4), 847–867. 10.1023/A:1024632923180

Zajonc, R. B. (1965). Social Facilitation: A solution is suggested for an old unresolved social psychological problem. Science, 149(3681), 269–274.

