## Supplemental material for "Social Tolerance and Innovation in Capuchins: socially more tolerant brown capuchins are better problem-solvers than less tolerant white-faced capuchins"

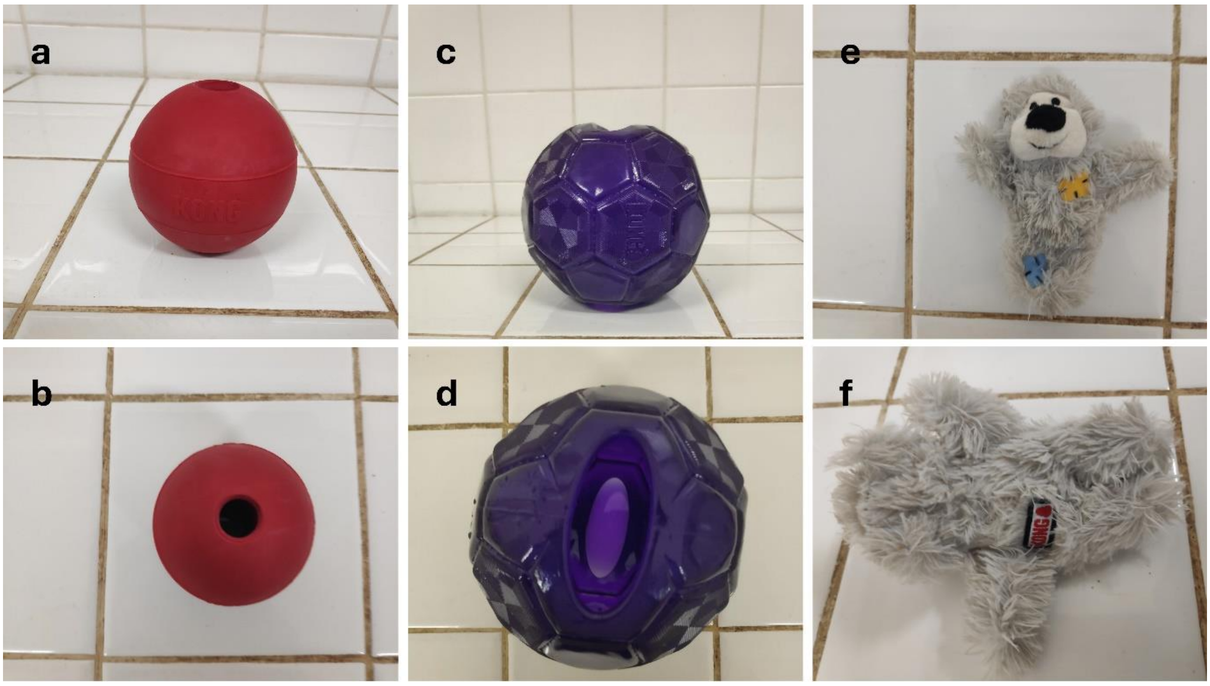


**Figure S1.** Novel objects used to test neophobia in white-faced capuchins (*Cebus capucinus*) and brown capuchins (*Sapajus apella*): a-b show a red ball with a small opening that could only be explored with a single finger; c-d show a purple ball with a larger opening, allowing individuals to insert a hand; e-f show a teddy-like plush toy. The red a purple balls were thoroughly cleaned before being presented to the next group. For sanitary reasons, cleaning was not feasible for the plush toy, so each group received a different novel plush toy of the same type.

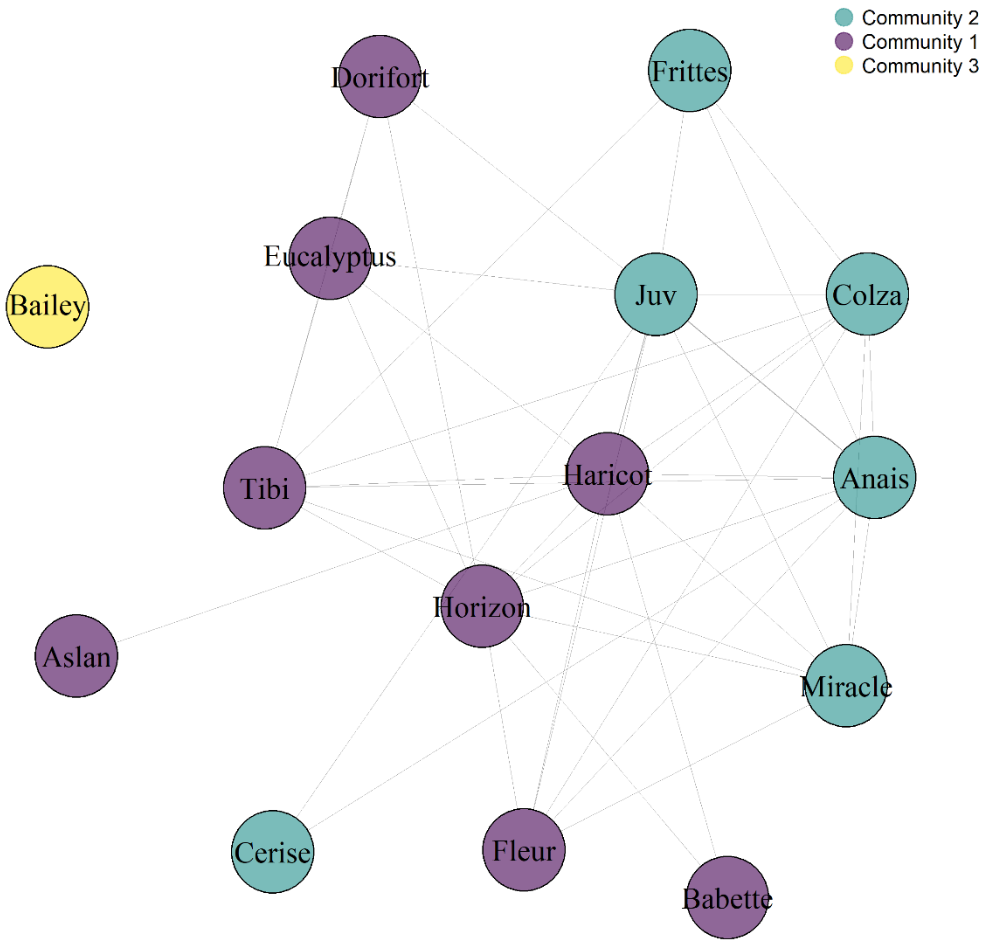


**Figure S2.** Social network of the large group of white-faced capuchins (*Cebus capucinus*) based on simple ratio associations (SRA) from the co-feeding experiment. Nodes represent individual capuchins, and edges indicate the strength of co-feeding associations, with thicker edges representing higher SRA values. Note, that overall the strength of associations are rather weak. The network reveals three distinct sub-communities within the group, suggesting non-random social structuring during feeding contexts. Community detection was based on modularity clustering.

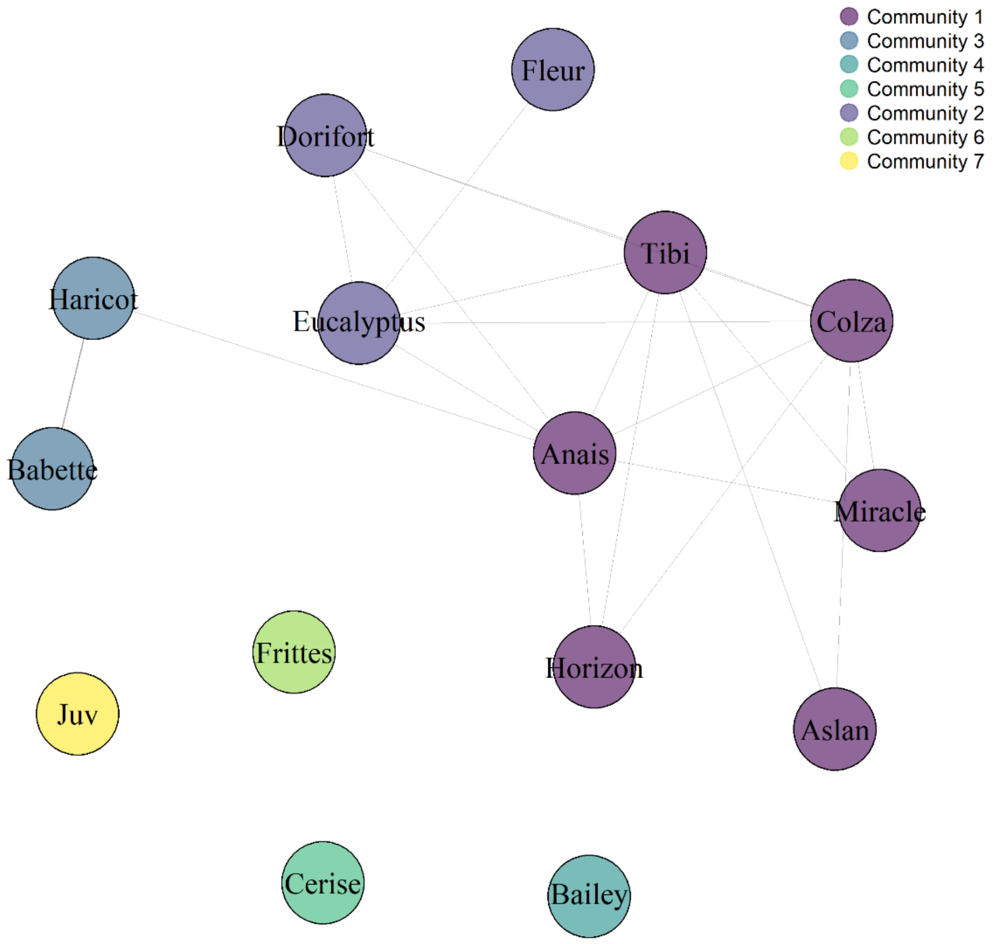


**Figure S3.** Social network of the large group of white-faced capuchins (*Cebus capucinus*) based on simple ratio associations (SRA) from the extractive foraging experiment. Nodes represent individual capuchins, and edges indicate the strength of associations during problem-solving, with thicker edges representing higher SRA values. Note, that overall the strength of associations are rather weak. The network reveals seven distinct sub-communities within the group, suggesting non-random social structuring during problem-solving contexts (four individuals that were not observed during the experiment and could potentially build one or multiple communities). Community detection was based on modularity clustering.

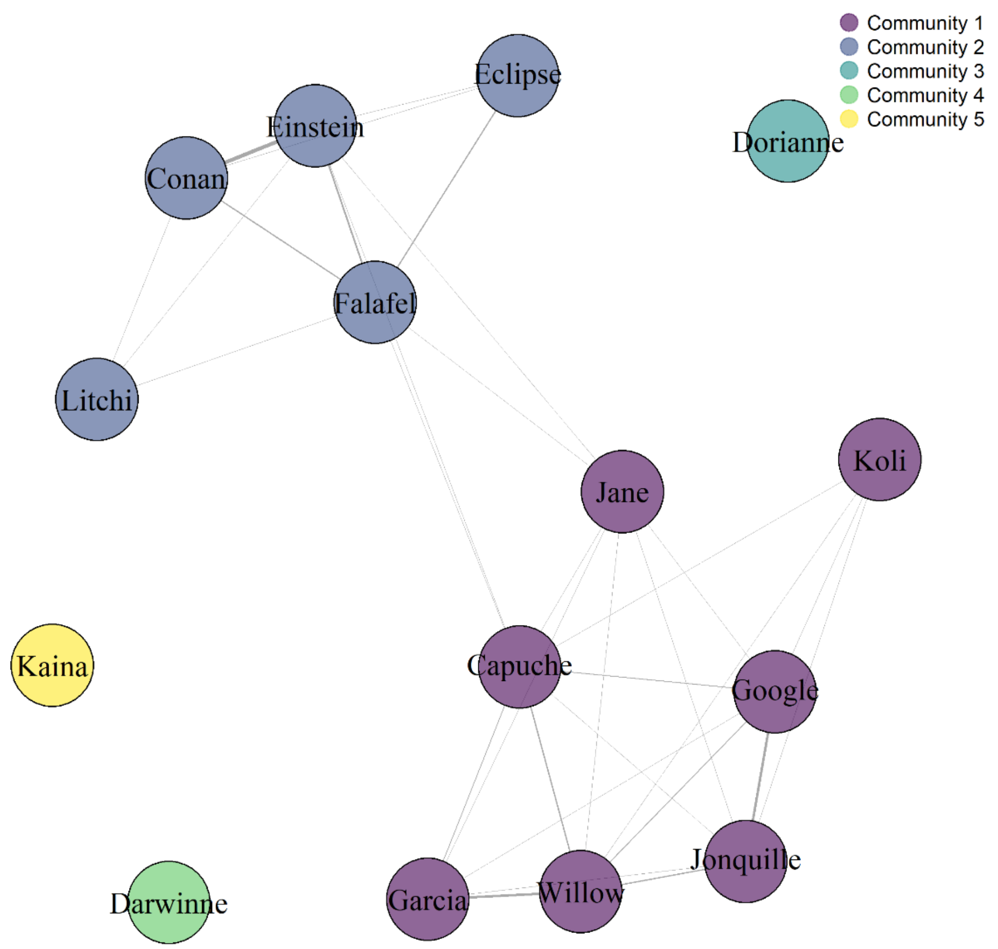


**Figure S4**. Social network of the large group of brown capuchins (*Sapajus apella*) based on simple ratio associations (SRA) from the co-feeding experiment. Nodes represent individual capuchins, and edges indicate the strength of co-feeding associations, with thicker edges representing higher SRA values. Note, that overall the strength of associations are stronger than in the white-faced capuchins. The network reveals five distinct sub-communities within the group, suggesting non-random social structuring during feeding contexts (four individuals that were not observed during the experiment and could potentially build one or multiple communities). Community detection was based on modularity clustering.

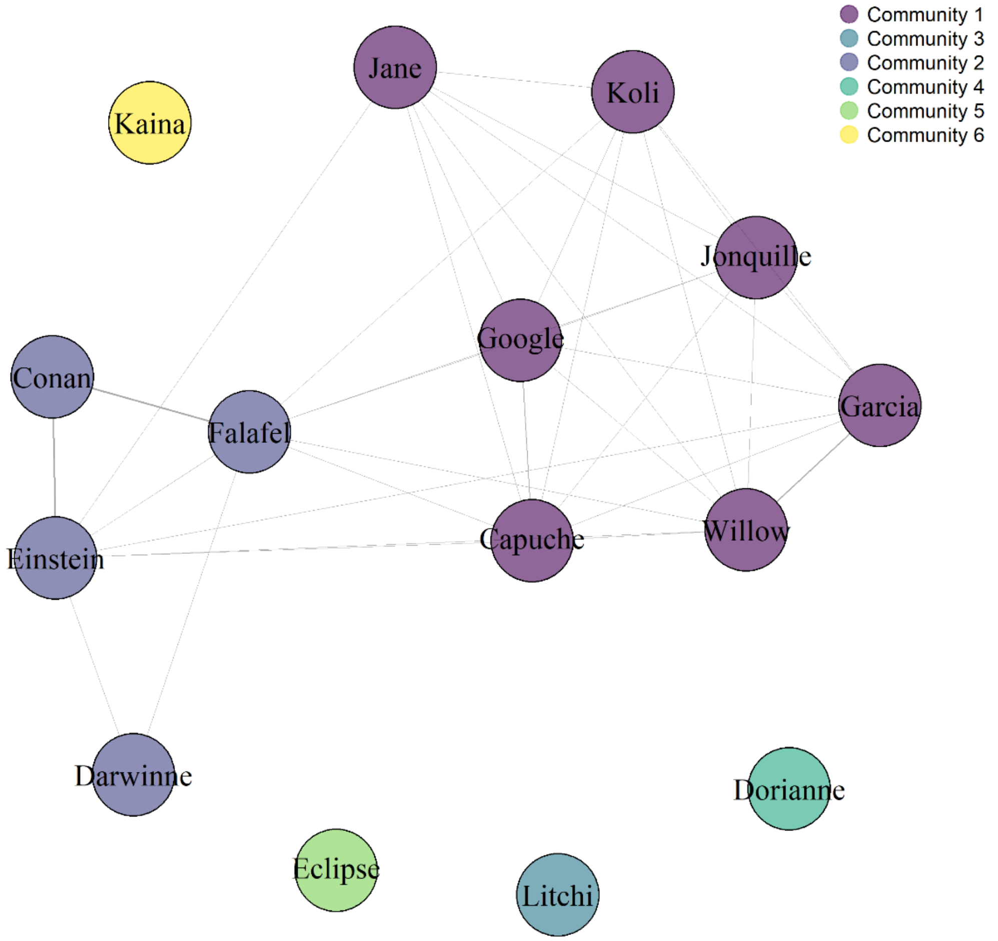


**Figure S5**. Social network of the large group of brown capuchins (*Sapajus apella*) based on simple ratio associations (SRA) from the extractive foraging experiment. Nodes represent individual capuchins, and edges indicate the strength of associations during problem-solving, with thicker edges representing higher SRA values. Note, that overall the strength of associations are stronger than in the white-faced capuchins. The network reveals six distinct sub-communities within the group , suggesting non-random social structuring during problem-solving contexts (four individuals that were not observed during the experiment and could potentially build one or multiple communities). Community detection was based on modularity clustering.

**Table S1.** Ethogram of the observed and analysed behaviours of the co-feeding and the food puzzle experiments.

| **Behaviour** | **Definition** |
|---|---|
| **Co-feeding** | |
| in the co-feeding area | Individuals were considered to be within the co-feeding area if at least two limbs were positioned on or within the imaginary lines extending from the markers at the corners of the area. Individuals sitting directly on top of the markers were also counted as inside the area, as they could still reach some of the food distributed within. In contrast, animals that only extended one arm into the area to retrieve individual pieces of apple were not considered to be within the co-feeding area. These instances were extremely rare, as the apple pieces were primarily distributed near the center of the area. |
| **Food puzzles** | |
| approaching | Individuals were considered to be within the food puzzle area if at least two limbs were positioned on or within the imaginary lines extending from the markers at the corners of the area. Individuals sitting directly on top of the markers were also counted as being inside the area. Although they could not access the food puzzles from that position, we assumed the proximity was sufficient for them to observe and potentially learn about the puzzle mechanics through social observation. |
| touching | Touching was defined as an individual either sitting on the puzzle or placing at least one hand on it without actively manipulating it. That is, the hand was resting passively on the puzzle without purposeful interaction. |
| sniffing | Sniffing was defined as the animal positioning its face in close proximity to the food puzzle, typically within a few centimeters, without necessarily making direct contact. |

| | |
|---|---|
| interacting | Interacting was defined as an individual actively moving its hand to manipulate the puzzle box in a goal-directed manner. |
| exploring | Individuals were classified as exploring the food puzzle if they engaged in behaviors such as sniffing, touching, or manipulating the puzzle in a goal-directed manner. Since the three behaviors (sniffing, touching, and interacting) were grouped into a single category of exploration, we established a clear priority system to avoid overestimating exploration when multiple behaviors occurred at the same time. Interaction was given the highest priority, followed by touching, and finally sniffing. For example, if an individual sniffed, touched, and interacted with the puzzle simultaneously, only the interaction was recorded. Similarly, if both sniffing and touching occurred, only sniffing was coded. This approach ensured that each instance of exploration was counted only once, based on the most active behavior observed. |
| solving | Solving was defined as an individual successfully extracting a food item from the food puzzle. |

**Table S2.** Summary of the GLMM for *proportion of individuals within the co-feeding area and food puzzle area per scan sample* (Model 1). Brown capuchins spent more time in the co-feeding and food puzzle areas than white-faced capuchins. Tolerance was reduced in the food puzzle condition and overall less individuals were present in the areas with increasing time.

| **fixed factor** | **estimate** | **SE** | **z** | **p** |
|---|---|---|---|---|
| Intercept | -2.43 | 0.41 | -5.78 | < 0.001 |
| species (*Sapajus apella*) | 1.84 | 0.59 | 3.13 | < 0.01 |
| condition (food puzzle) | -0.49 | 0.16 | -2.98 | < 0.01 |
| z-transformed scan | -0.24 | 0.11 | -2.14 | 0.03 |

**Table S3.** Summary of the mixed effects Cox proportional hazards model for the latency of approaching the food puzzles. Significant predictors are indicated with a bold proportional hazards ratio. For the influence of the task, the linear relationship is depicted.

| **fixed factor** | **proportional hazards ratio** | **95% CI** | **SE** | **z** |
|---|---|---|---|---|
| species (*S. apella*) | 2.11 | 0.83 – 5.34 | 0.47 | 1.57 |
| sex (male) | 1.36 | 0.71 – 2.63 | 0.33 | 0.93 |
| age | 1.09 | 1.03 – 1.14 | 0.03 | 3.25 |
| task (L) | 0.91 | 0.62 – 1.35 | 0.2 | -0.45 |

**Table S4.** Summary of the GLMM for the number of approaches. Significant predictors are indicated with a bold odds ratio (OR). For the influence of the task, the linear relationship is depicted.

| **fixed factor** | **OR** | **95% CI** | **estimate ± (SE)** | **z** |
|---|---|---|---|---|
| Intercept | 0.49 | 0.10 – 2.39 | -0.69 ± 0.8 | -0.87 |
| species (*S. apella*) | 0.96 | 0.26 – 3.58 | -0.04 ± 0.67 | -0.06 |
| sex (male) | **7.38** | 2.65 – 20.54 | 1.99 ± 0.52 | 3.83 |
| age | **1.13** | 1.05 – 1.23 | 0.13 ± 0.04 | 3.19 |
| task (L) | **1.17** | 1.07 – 1.28 | 0.16 ± 0.05 | 3.41 |

**Table S5.** Summary of the mixed effects Cox proportional hazards model for the latency of exploration the food puzzles. Significant predictors are indicated with a bold proportional hazards ratio. For the influence of the task, the quadratic relationship is depicted.

| **fixed factor** | **proportional hazards ratio** | **95% CI** | **SE** | **z** |
|---|---|---|---|---|
| species (*S. apella*) | **2.81** | 1.08 – 7.28 | 0.49 | 2.12 |
| sex (male) | 2.64 | 0.94 – 7.36 | 0.52 | 1.85 |
| age | **1.18** | 1.09 – 1.28 | 0.22 | 3.99 |
| task (Q) | **0.55** | 0.35 – 0.87 | 0.23 | -2.56 |

**Table S6.** Summary of the GLMM for the number of explorative actions. Significant predictors are indicated with a bold odds ratio (OR). For the influence of the task, the quadratic relationship is depicted.

| **Fixed factor** | **OR** | **95% CI** | **Estimate ± (SE)** | **z** |
|---|---|---|---|---|
| Intercept | **0.05** | 0.01 – 0.49 | -2.93 ± 1.13 | -2.58 |
| species (*S. apella*) | 4.52 | 0.75 – 27.23 | 1.51 ± 0.91 | 1.65 |
| sex (male) | **18.11** | 4.17 – 78.72 | 2.89 ± 0.75 | 3.86 |
| age | **1.28** | 1.14 – 1.44 | 0.25 ± 0.06+ | 4.18 |
| task (Q) | **0.84** | 0.81 – 0.88 | -0.17 ± 0.02 | -8.15 |

**Table S7.** Summary of the mixed effects Cox proportional hazards model for the latency of successful extractions of food. Significant predictors are indicated with a bold proportional hazards ratio. For the influence of the task, the quadratic relationship is depicted.

| **fixed factor** | **proportional hazards ratio** | **95% CI** | **SE** | **z** |
|---|---|---|---|---|
| species (*S. apella*) | **4.08** | 1.24 – 13.39 | 0.61 | 2.32 |
| sex (male) | 1.51 | 0.62 – 3.72 | 0.46 | 0.9 |
| age | **1.23** | 1.12 – 1.34 | 0.05 | 4.54 |
| task (Q) | 0.79 | 0.38 – 1.65 | 0.38 | -0.63 |

**Table S8.** Summary of the GLMM for the proportion of successful extractions. Significant predictors are indicated with a bold odds ratio (OR). For the influence of the task, the quadratic relationship is depicted.

| **Fixed factor** | **OR** | **95% CI** | **Estimate ± (SE)** | **z** |
|---|---|---|---|---|
| Intercept | **4.5e-08** | 3.7e-10 – 5.5e-06 | -16.9 ± 2.45 | -6.9 |
| species (*S. apella*) | **555.8** | 29.7 – 10404.9 | 6.32 ± 1.49 | 4.23 |
| sex (male) | **23.9** | 1.40 – 407.2 | 2.89 ± 0.75 | 3.86 |
| age | **1.28** | 1.14 – 1.44 | 3.17 ± 1.45+ | 2.2 |
| task (Q) | **5.52** | 4.67 – 6.55 | 1.71 ± 0.09 | 19.77 |

**Table S9.** Summary of the GLMM for the proportion of successful extractions for brown capuchins. Significant predictors are indicated in bold.

| fixed factor | estimate | SE | z | p |
|---|---|---|---|---|
| **Intercept** | **-8.11** | **1.28** | **-6.31** | **< 0.001** |
| neophobia | -0.22 | 0.76 | -0.29 | 0.77 |
| eigenvector centrality (co-feeding) | -2.85 | 5.77 | -0.5 | 0.62 |
| eigenvector centrality (food puzzle) | 3.25 | 6.61 | 0.49 | 0.62 |
| sex (male) | 0.48 | 1.7 | 0.28 | 0.78 |
| **age** | **0.39** | **0.1** | **3.83** | **< 0.001** |
